# Phospho-proteomic analysis of CAR-T cell signaling following activation by antigen-presenting cancer cells

**DOI:** 10.1101/2022.02.24.481820

**Authors:** Melanie A. MacMullan, Zachary S. Dunn, Yun Qu, Pin Wang, Nicholas A. Graham

## Abstract

Chimeric antigen receptors (CARs) are synthetic biomolecules comprised of an extracellular antigen recognition domain and intracellular signaling domains. When expressed in immune cells, CARs direct their host cells to kill diseased cells expressing the antigen recognized by the CAR. Although signaling pathways downstream of CAR activation control the cytotoxic function of CAR-expressing cells, phospho-proteomic studies of CAR signaling have been limited. Most approaches have used antibodies or soluble ligands, rather than cell-displayed antigens, to activate CAR signaling. Here, we demonstrate an efficient and cost-effective label-free phospho-proteomic approach to analyze CAR signaling in immune cells stimulated with antigen-presenting cancer cells. Following co-culture of CAR-T cells with cancer cells, we first preserve phospho-signaling by cross-linking proteins with formalin. Then, we use magnet-activated cell sorting (MACS) to isolate CAR-T cells from the co-culture. Validation experiments demonstrated that formalin fixation did not alter the phospho-proteome and that MACS achieved >90% CAR-T cell purity. Next, we compared the phospho-proteome in CAR-T cells stimulated with either CD19-expressing or non-CD19-expressing SKOV3 ovarian cancer cells. This analysis revealed that CAR signaling activated known pathways including the mitogen-activated protein kinases (MAPKs) ERK1/2. Bioinformatic approaches further showed that CAR activation induced other signaling pathways including the MAPK p38α, protein kinase A, and checkpoint kinase 1 (CHK1). Taken together, this work presents an easy and inexpensive method to better understand CAR immunotherapy by label-free phospho-proteomic analysis of CAR signaling in immune cells stimulated by antigen-presenting cancer cells.

## Introduction

Immunotherapy harnesses components of the immune system to fight cancer and other diseases. Immune cells engineered to express chimeric antigen receptors (CARs) are one of the most promising forms of immunotherapy (1). CARs are synthetic biomolecules comprised of modular protein domains including a ligand-binding domain (e.g., single-chain variable fragment (scFv)), an extracellular spacer domain, a transmembrane domain, and intracellular signaling/co-stimulation domains. When expressed in T cells, CARs repurpose endogenous T cell signaling pathways to kill cancer cells that express a target recognized by the scFv. CAR-T cells have shown great therapeutic promise, and several CAR-T cell therapies have gained FDA approval (2–4). However, despite extensive research into CAR-T cell immunotherapy, there remain significant hurdles for CAR-T cells to become a “standard of care” therapy including improving efficacy and persistence while minimizing adverse events such as neurotoxicity and cytokine release syndrome (5). Thus, more studies are urgently needed to better understand and optimize CARs to improve CAR immunotherapy.

Because CAR signaling controls the cytotoxic function of T cells, the design of next-generation CARs that overcome current limitations would benefit from a rigorous and comprehensive understanding of CAR signaling (6). Because CARs are built from endogenous proteins involved in T cell receptor (TCR) signaling, it has been thought that their activation pathways share many similarities. However, emerging data suggests that CARs activate a mixture of canonical TCR signaling and unique CAR-specific pathways (7–10). In addition, the choice of CAR stimulatory domains (e.g., CD28, 4-1BB, or both) can influence the strength of CAR signaling, thereby affecting CAR-T cell function (11–13). Therefore, more research is needed to generate a quantitative understanding of CAR signaling and how it controls the cytotoxic function of CAR-T cells.

For comprehensive, unbiased, and quantitative characterization of protein phosphorylation, liquid chromatography-mass spectrometry (LC-MS)-based phospho- proteomics is the technology of choice (14), and these methods have been applied to CAR signaling (7, 9, 11, 15). However, phospho-proteomic studies of CAR signaling have been limited by the challenge of stimulating CAR-expressing cells with their *in vivo* stimulus, antigen-presenting cancer cells. In LC-MS-based proteomic studies, mixing two cell types such as CAR-T and target cancer cells confounds analysis because both cell types express many of the same proteins. Additionally, because phospho-signaling occurs on rapid time scales (16, 17), it is not feasible to use flow sorting to separate cell types before phospho-proteomic analysis. As such, some studies have relied on antigen-presenting beads to activate CARs (9, 11, 18) but these methods fail to produce a realistic immunological synapse with membrane interaction and ligand exchange. To overcome this limitation, others have used stable isotope labeling with amino acids in cell culture (SILAC) to label CAR-T cells and cancer cells before co-culture, thereby enabling deconvolution of peptides on the MS (7, 15). However, SILAC-based methods remain expensive and limited in the number of samples that can be quantified and the number of cell types that can be co-cultured (19, 20).

To overcome these limitations, we have developed a formalin fixation and cell sorting method that enables label-free phospho-proteomic analysis of CAR-T cells stimulated with antigen-expressing cancer cells. Specifically, after mixing CAR-T and cancer cells, we preserve transient phospho-signals by formalin fixation, label CAR-T cells with magnetic beads coated with an antibody that recognizes the CAR, and then purify CAR-T cells using magnet-activated cell sorting (MACS) (21) (Fig. 1). Isolated CAR-T cells are then lysed, and formalin-fixed proteins are de-crosslinked and analyzed by phospho-proteomics. The method presented here represents an easy and inexpensive method for phospho-proteomic analysis of CAR signaling in immune cells stimulated by cancer cells expressing the target antigen.

**Figure 1.**
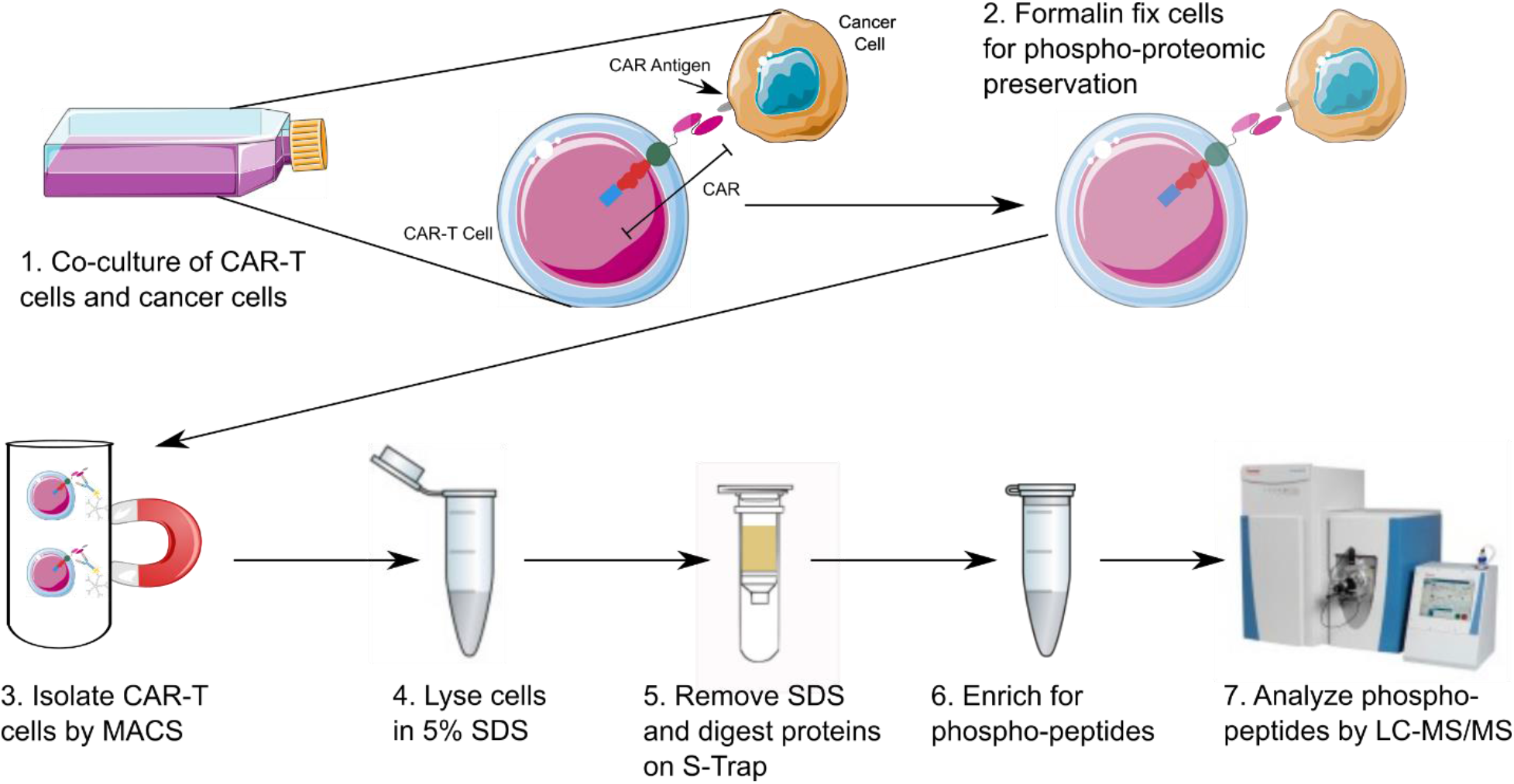
Workflow for phospho-proteomic analysis of CAR signaling in CAR-T cells stimulated with antigen-expressing cancer cells. To activate CAR signaling, CAR-T cells are co-cultured with antigen-expressing cancer cells. To preserve protein phosphorylation, the cell mixture is then fixed with formalin. Next, CAR-T cells are separated from cancer cells by MACS after labeling with a biotinylated antibody that recognizes the CAR and streptavidin-coated magnetic Dynabeads. The MACS eluate is then lysed and formalin crosslinks are reversed using an SDS lysis buffer. Using an S- trap, SDS is removed from the lysate, and proteins are digested to peptides with trypsin. Lastly, phospho-peptides are purified with TiO2 beads and analyzed by LC-MS.

## Materials and Methods

### Plasmid Construction

The retroviral vector encoding anti-CD19 CAR was constructed by incorporating the anti-CD19 ScFv derived from the anti-CD19 antibody FMC63 (Acro Biosystems) into the MP-71 retroviral vector backbone kindly provided by Prof. Wolfgang Uckert (Max-Delbrück-Center for Molecular Medicine) using methods previously described (22). The CAR expression cassette within MP-71 backbone also contains a CD8 hinge, CD8 and CD3ζ transmembrane domains, and CD28 and CD3ζ signaling domains. The lentiviral vectors encoding GFP and CD19 were constructed from the FUW backbone as previously described (23).

### Vector Production

293T cells were obtained from ATCC (CRL-3216) and maintained in D10 medium consisting of DMEM supplemented with 10% fetal bovine serum (FBS), 2 mM L-glutamine, and 0.35% Pen Strep (Gibco 15140-122). Media was filtered using a 0.2-μm bottle filter and warmed to 37°C before being added to cells. Retroviral and lentiviral vectors were prepared by transient transfection of 293T cells using a standard calcium phosphate precipitation method. Briefly, 18 million 293T cells were seeded in a 15-cm tissue-culture dish. When confluency reached 70%–80%, retrovirus was produced by transfecting 293T cells with plasmids encoding the retroviral backbone, the RD114 envelope protein and the packaging proteins (Gag-Pol). For lentivirus, 293T cells were transfected with plasmids encoding the lentiviral backbone, VSV-G envelope protein, and packaging proteins (Gag-Pol). The viral supernatants were harvested 48 hours after transfection and filtered through a 0.45-μm filter (Corning). Filtered viral supernatant was ultracentrifuged at 25,000 rpm for 1.5 hours to increase viral titer.

### Anti-CD19 CAR-T Cell Production

Frozen human peripheral blood mononuclear cells (PBMCs) (AllCells by FisherScientific, 50-156-486) and were recovered in T-cell culture medium (TCM) consisting of AIM-V Medium (ThermoFisher, 12055083), 5% human AB serum (GemCell, GeminiBio, 100-512), 10 mM HEPES, 1% GlutaMax-100x, 12.25 mM N-acetylcysteine, and 0.35% Pen Strep (Gibco, 15140-122). Media was filtered using a 0.2-μm bottle filter and warmed to 37°C before being added to cells. Interleukin-2 (IL-2) (PeproTech, 200-02) was added at a concentration of 100 U/mL during each media refresh. PBMCs were recovered at a concentration of 1 × 10^6^ cells/mL overnight in a 5% CO2, 37°C and humidified incubator. T cells were activated by adding Dynabeads Human T-Expander CD3/CD28 (Invitrogen, 11141D) at a bead:PBMC ratio of 3:1 for 48 hours in a 5% CO2, 37°C and humidified incubator. For T cell transduction, RetroNectin (TaKara, T202) was bound to a non-tissue culture treated plate overnight at 4°C. Both T cells and viral supernatants were added to the RetroNectin and centrifuged for 90 minutes at 300 g. Fresh media was added to the cells and transduced T cells were expanded for 2 weeks in TCM, during which time the culture medium was replenished every 2 days and T cell density was maintained between 0.5 and 1 × 10^6^ cells/mL. Following expansion, T cells were then reconstituted in freezing medium (90% FBS, 10% DMSO) and stored in the vapor phase of a liquid nitrogen cell stock dewar until needed.

### Cell Culture

Anti-CD19 CAR-T cells were expanded in TCM. Interleukin-2 (IL-2) was added at a concentration of 100 U/mL during each media refresh. Cells were initially recovered at a concentration of 1 × 10^6^ cells/mL but maintained at a concentration between 0.5 and 1 × 10^6^ cells/mL for all subsequent passages. SKOV3 cells were obtained from ATCC (30–2007) and transduced with either the lentiviral vector FUW-CD19 or the lentiviral vector FUW-GFP to overexpress CD19 or GFP, respectively. Non-transduced SKOV3 cells were also used as controls in these experiments. SKOV3 cells were expanded in RPMI media containing 15% FBS (Sigma, F2442), 2 mM L-glutamine (Corning, MT25005CV), 20 mM HEPES (ThermoFisher Scientific, 15630106), 1 mM β-mercaptoethanol (Gibco, 21985- 023), and 0.35% Pen Strep (Gibco, 15140-122). All cells were counted by Trypan Blue exclusion with a hemocytometer. All cells were grown in a 5% CO2, 37°C and humidified incubator and were used within 20 passages of thawing.

### Flow Cytometry Analysis

To measure interferon gamma (IFNγ) production, cells were mixed at effector-to-target ratios of 10:1, 5:1, 1:1, 0.5:1, and 0.2:1 to determine which ratio prompted the greatest response. Cells were treated with 5 µg/mL of Brefeldin A (BioLegend, 420601) and allowed to incubate for 6 hours in a 5% CO2, 37°C and humidified incubator. Cells were then washed 1 time with 100 µL PBS and then fixed and permeabilized in 100 µL Fixation and Permeabilization Solution from the BD Cytofix/Cytoperm Fixation/Permeabilization Kit (BD BioSciences, 554714) for 10 minutes at 4°C. Cells were washed 2 times using 100 µL 1X Perm/Wash buffer provided in the Cytofix/Cytoperm Fixation/Permeabilization Kit and then labeled with a R-Phycoerythrin (PE) labeled anti-human interferon(IFN)-gamma(γ) antibody at a ratio of 1:100 in 100 µL 1X Perm/Wash buffer and incubated for 30 minutes at 4°C. Cells were then washed 2 times using 100 µL 1X Perm/Wash buffer and resuspended in 100 µL PBS for flow cytometry analysis. Cells were then analyzed using a Miltenyi Biotec MACSQuant benchtop flow cytometer.

To measure cytotoxicity, SKOV3 target cells were resuspended in 0.1% BSA in PBS to a concentration of 1 × 10^6^ cells/mL and stained with 1 µM CFSE for 10 min at 37°C. 1 equal volume of FBS was added to stop the reaction, and cells were washed 3X with PBS to remove excess CFSE before being reconstituted in TCM. Cells were then mixed at a target-to-effector ratio of 1:1 and allowed to incubate for 24 hours in a 5% CO2, 37°C and humidified incubator. Cells were then washed 1 time with 100 µL PBS and stained with 7AAD for 10 minutes at 4°C. Cells were then analyzed using a Miltenyi Biotec MACSQuant benchtop flow cytometer.

### Formalin Fixation Experiments

T cells expanded from the same PBMC donor as other experiments used in this study were evaluated for the effect of protein crosslinking by formalin fixation on phospho- proteomic results. The non-fixed population (NFF) was briefly centrifuged and washed once with PBS before being immediately frozen at -80°C while the experimental population was fixed (FF) by adding an equal volume of 20% neutral buffered formalin for a final formaldehyde concentration of ∼4%. Both FF and NFF populations were lysed according to the same protocol in a 5% SDS lysis buffer as described below. Three biological replicates of each condition were collected and analyzed.

### Cell Co-culture and Magnet-associated Cell Sorting (MACS)

Co-cultures of CAR-T cells and SKOV3.CD19 or SKOV3.NT were incubated at a 1:1 target-to-effector ratio for 45 minutes in a 5% CO2, 37°C and humidified incubator. Immediately following co-culture, cells were mixed with 20% neutral buffered formalin for a final formaldehyde concentration of ∼4%. For analysis of MACS purity, SKOV3.NT and SKOV3.CD19 cells were first labeled with either CFSE or GFP. CFSE labeling was done as described for the cytotoxicity assay. GFP labeling was performed via transfection with the FUW-GFP vector. Labeled SKOV3.NT and SKOV3.CD19 cells were mixed with CAR-T cells for 45 min and then formalin fixed as described above. Next, the cell mixture was washed with PBS, and CD19.CAR-T cells were labeled with biotinylated anti-Fab antibodies (ThermoFisher Scientific, 31803). The co-culture was then mixed with CELLection Biotin Binder (ThermoFisher Scientific, 11533D) Dynabeads for 20 minutes at 4°C with rotation. A magnetic rack was used to pull Dynabead bounds cells out of solution, and the cell-bound beads were washed 2x with 0.1% BSA in PBS and then treated with a DNase I-containing release buffer provided by the CELLection kit to break the DNA linker and release the cells from the beads. Isolated cells were then analyzed in technical duplicate by LC-MS proteomics or flow cytometry for CFSE or GFP signal to evaluate purity.

### Cell Lysis and Formaldehyde Removal

To de-cross link proteins following formalin fixation, cells were sonicated in a 5% SDS, 100 mM Tris buffer and then heated for 60 minutes at 80°C (24). This process was repeated once more before proteins were reduced with 10 mM dithiothreitol (DTT) for 30 minutes at room temperature (RT), alkylated with 20 mM iodacetamide (IAA) for 30 minutes at RT in the dark, and quenched with 10 mM DTT. Samples were then diluted 8x with S-Trap buffer (90% methanol, 100 mM triethylammonium bicarbonate (TEAB) (ThermoFisher, 90114)) solution. The protein solution was then added directly to S-Trap mini columns (Protifi, C02-mini-40) by centrifugation (400 µL at a time, 4,000 g spin for 30 seconds) and washed 3x with S-Trap buffer to remove SDS (25). 125 µL per column of Trypsin solution was prepared by diluting Trypsin (Pierce Trypsin Protease, 90058) at a 1:20 protein mass ratio in digestion buffer (50mM TEAB). Proteins were digested overnight in a 37°C water bath. The following morning, the columns were placed in new collections. 80 µL of digestion buffer was added to each column, and centrifugation for 1 minute at 1000x g was used to collect the elution. 80 µL of 0.2% formic acid (FA) was added to the column and elution was collected into the same tube following centrifugation for 1 minute at 1000x g. A final 80 µL of 50% acetonitrile (ACN) / 0.2% FA was added to the column and the elution was collected into the same tube following centrifugation for 1 minute at 1,000 g. Peptides were then vacuum dried prior to phospho-enrichment or directly injected onto the LC-MS for proteomic analysis.

### LC-MS Phospho-proteomics

300 µg of dried peptides were resuspended in 170 µL loading buffer (65% ACN, 2% TFA, saturated with 8.6 mg/mL glutamic acid) for phospho-enrichment. Titanium dioxide beads (GL sciences, #1400B500) were weighed out at a ratio of 10 mg beads to 1 mg tryptic peptide for enrichment and resuspended in 30 µL per sample of loading buffer to prepare a bead slurry. Samples and beads were vortexed (setting 6.0 on Vortex-Genie 2T, SIIE10013) for 15 minutes at room temperature (RT) to ensure mixing. 30 µL of bead slurry was aliquoted into a new tube for each sample. Reconstituted dried peptides were added to the bead aliquots and samples were vortexed (setting 6.0 on Vortex-Genie 2T) at RT for 60 minutes. Samples were centrifuged and the supernatant was discarded. 200 µL of loading buffer was added to each sample and vortexed (setting 6.0 on Vortex-Genie 2T) at RT for 30 minutes followed by centrifugation and discard of the supernatant. Samples were resuspended in 200 µL wash buffer 1 (WB1, 65% ACN, 0.5% TFA) and vortexed (setting 6.0 on Vortex-Genie 2T) at RT for 30 minutes followed by centrifugation and discard of the supernatant. Samples were resuspended in 200 µL wash buffer 2 (WB2, 65% ACN, 0.1% TFA) and vortexed (setting 6.0 on Vortex-Genie 2T) at RT for 30 minutes followed by centrifugation and discard of the supernatant. Samples were resuspended in 200 µL elution buffer 1 (EB1, 50% ACN, 0.3M NH4OH) and incubated for 1 hour on a ThermoMixer at 45°C, 1,400 RPM. Following centrifugation, the supernatant was transferred to a clean tube and the tube was placed in a speed vac to begin the drying down process. Beads were then reconstituted in 200 µL elution buffer 2 (EB2, 5% ACN, 0.3M NH4OH) and incubated for 1 hour on a ThermoMixer at 45°C, 1,400 RPM. Following centrifugation, the supernatant was transferred to their corresponding tubes and samples were dried down.

Desalting was carried out on stage tips composed of empty pipette tips loaded with two cores of C18. C18 stage tips were conditioned with two times 150 µL of 50% ACN followed by two times 150 µL of 0.1% TFA. Reagents were passed through the stage tips by centrifugation (2,000-3,000 g). Samples were resuspended in 100 µL of 0.1% TFA and transferred onto the stage tips. Stage tips were placed in the original sample tubes to effectively “wash” the tube as the sample eluted. Elutions were transferred to the stage tip for another passage. Stage tips were then washed 3 times with 150 µL 0.1% TFA. Samples were eluted with 75 µL of 60% ACN, 0.1%TFA and dried down to completion. Dried phospho-peptides were resuspended in 6 µL of LC buffer A (0.1% FA).

The samples were randomized and 5 µL of each sample was injected into an Easy 1200 nanoLC ultra high-performance liquid chromatography coupled with a Q Exactive Plus quadrupole-Orbitrap mass spectrometer (Thermo Fisher Scientific). Peptides were separated by a reverse-phase analytical column (PepMap RSLC C18, 2 µm, 100 Å, 75 µm×25 cm). Flow rate was set to 300 nL/min at a gradient from 3% LC buffer B (0.1% formic acid, 80% acetonitrile) to 38% LC buffer B in 110 min, followed by a 10-min washing step to 85% LC buffer B. The maximum pressure was set to 1,180 bar, and column temperature was maintained at 50°C. Peptides separated by the column were ionized at 2.4 kV in positive ion mode. MS1 survey scans were acquired at the resolution of 70,000 from 350 to 1,800 m/z, with a maximum injection time of 100 ms and AGC target of 1e6. MS/MS fragmentation of the 14 most abundant ions were analyzed at a resolution of 17,500, AGC target 5e4, maximum injection time 65 ms, and normalized collision energy of 26. Dynamic exclusion was set to 30 sec, and ions with charge +1, +7 and >+7 were excluded.

MS/MS fragmentation spectra were searched with Proteome Discoverer SEQUEST (version 2.2, Thermo Scientific) against in silico tryptic digested Uniprot all-reviewed Homo sapiens database (release August 2017, 42,140 entries) plus all recombinant protein sequences used in this study. The maximum missed cleavages was set to two. Dynamic modifications were set to phosphorylation on serine, threonine or tyrosine (+79.966 Da), oxidation on methionine (+15.995 Da), and acetylation on protein N-terminus (+42.011 Da). Carbamidomethylation on cysteine (+57.021 Da) was set as a fixed modification. The maximum parental mass error was set to 10 ppm, and the MS/MS mass tolerance was set to 0.02 Da. The false discovery threshold was set strictly to 0.01 using the Percolator Node validated by q-value. The relative abundance of parental peptides was calculated by integration of the area under the curve of the MS1 peaks using the Minora LFQ node. A signal-to-noise threshold of 1.5 was implemented. Individual phospho-site localization probabilities were determined by the ptmRS node, and phospho-sites with <0.75 localization probability were removed. Peak files were obtained through conversion by the MSConvertGUI (v 3.0.21301-4175933). RAW LC-MS files and processed result files are available at MassIVE database (26) under identifier MSV000088632 (username: MSV000088632_reviewer, password: MM2021).

### Experimental Design and Statistical Rationale

For phospho-proteomic analysis of CAR-T signaling, CAR-T cells were co-cultured with CD19-expressing SKOV3 cells (SKOV3.CD19) as the experimental condition or with non-CD19-expressing SKOV3 cells (SKOV3.NT) as the control. Three independent biological replicates were used for each condition. These samples were then run in technical duplicate on the LC-MS. The label-free phospho-peptide data was filtered for phospho-peptides having two or fewer missed cleavages. No data was imputed. Phospho-peptide intensities were log2 transformed and median normalized. To analyze the effect of formalin fixation on the phospho-proteome, phospho-peptide intensities and identities were compared directly. For analysis of the CAR-T signaling phospho-proteomic dataset, we applied strict filters to ensure that each phospho-peptide included in the analysis was 1) observed in at least one technical replicate of every sample and 2) observed in at least 10 of 12 replicates. We then performed an error propagation on the technical replicates to calculate the standard deviation of our biological replicates and used these values to perform a two-sample t-test. P-values were corrected for multiple hypotheses testing using the Benjamini-Hochberg method. Post-translational modification-signature enrichment analysis (PTM-SEA) was performed according to the established workflow (27).

## Results

### Anti-CD19 CAR-T cells kill CD19-expressing SKOV3 cells

Human T cells were expanded from PBMCs and then engineered to express a second-generation CAR that recognizes the tumor antigen CD19. Following recovery from freeze-thaw, 51% of the transduced T cells expressed the anti-CD19 CAR (Fig. 2A). We next tested whether stimulation with SKOV3 ovarian cancer cells expressing either CD19 (SKOV3.CD19) or not (SKOV3.NT) could stimulate CAR-T cells to produce IFNγ, a marker of activated T cells. We tested several effector-to-target (E:T) cell ratios and found that stimulation with SKOV3.CD19 cells but not SKOV3.NT cells induced IFNγ production in CAR-T cells (Fig. 2B). The maximal IFNγ response occurred with a 1:1 E:T cell ratio. We thus tested the extent of cell killing of CAR-T cells co-cultured with SKOV3.NT or SKOV3.CD19 cells at a 1:1 E:T cell ratio and found that anti-CD19 CAR-T cells mediated five-fold more killing of SKOV3.CD19 than SKOV3.NT (Fig. 2C). Taken together, this data demonstrates that anti-CD19 CAR-T cells respond to and kill cancer cells in a CD19-specific fashion.

**Figure 2.**
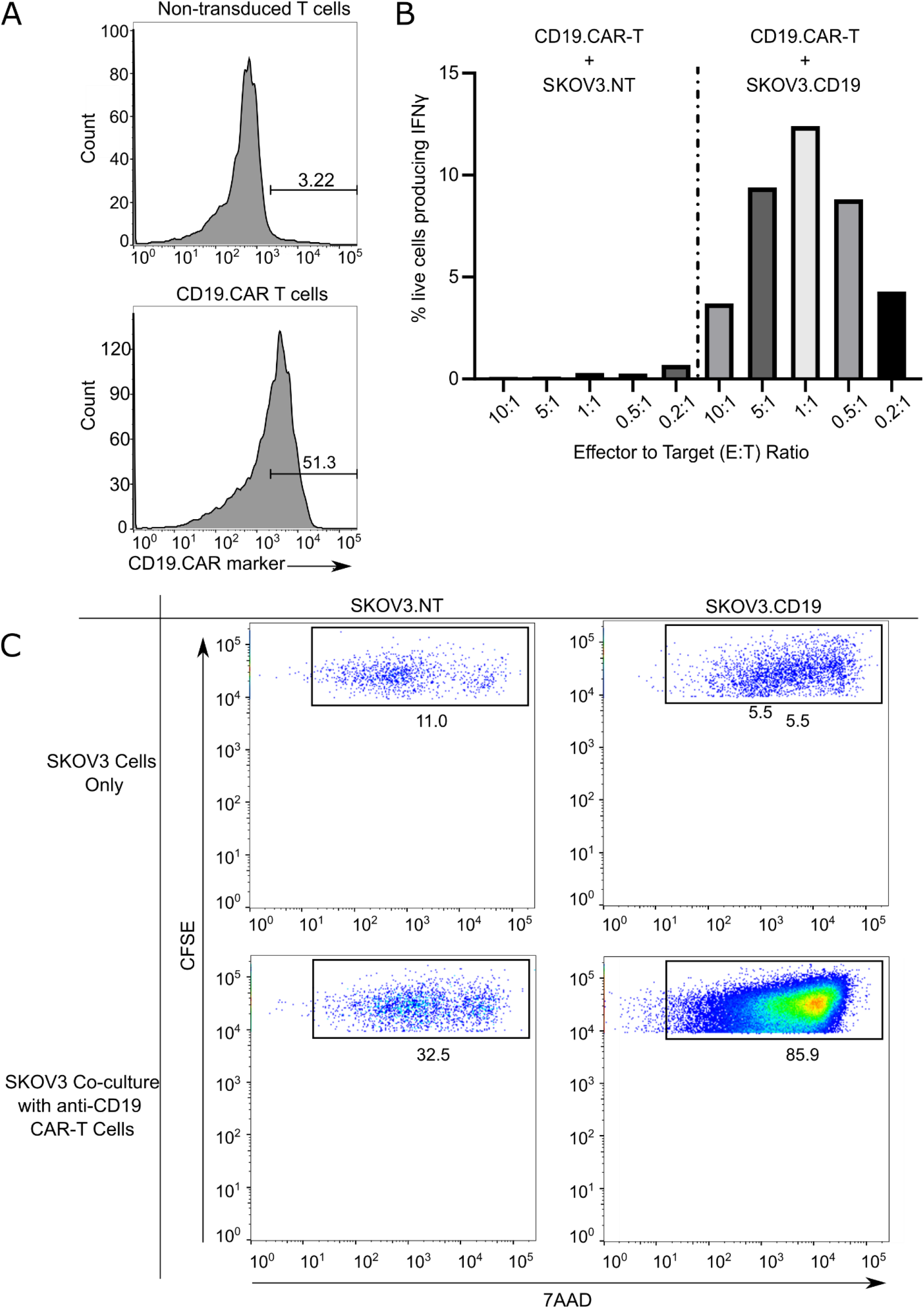
Anti-CD19 CAR-T cells kill CD19-expressing SKOV3 cells. A) Following expansion from human PBMCs, T cells were either not transduced or transduced with a retroviral vector encoding a second-generation CAR against CD19. Following freeze- thaw, flow cytometry with an antibody that recognizes the anti-Fab region of the CAR demonstrated that 48% of transduced T cells expressed the CAR. B) CAR-T cells were mixed with non-transduced SKOV3 cells (SKOV3.NT) or CD19-expressing SKOV3 cells (SKOV3.CD19) at the shown effector-to-target (E:T) cell ratios. Interferon (IFN)- gamma(γ) production was measured by flow cytometry 6 hours later. C) SKOV3.NT and SKOV3.CD19 cells were stained with CFSE and then mixed or not mixed with CAR-T cells at a 1:1 E:T ratio. 24 h later, the percentage of dead SKOV3 cells was measured by 7AAD staining using flow cytometry.

### Formalin fixation does not impact phospho-peptide identification or quantitation by LC-MS

Next, we tested the effects of formalin fixation on the identification and quantitation of phospho-peptides by LC-MS (Supp. Table 1). We prepared parallel samples of T cells either by formalin fixation or flash freezing, followed by lysis in 5% SDS buffer, and SDS removal and on-column protein digestion using S-traps (25). Phospho-peptides were then enriched by TiO2 pulldown and were analyzed by LC-MS. In both formalin-fixed and flash-frozen samples, we identified 5,254 phospho-peptides from 2,110 proteins, of which 4,569 phospho-peptides from 1,931 proteins were quantified in at least 1 of 3 replicates per condition with 93.2% of phospho-peptides identified in both conditions (Fig. 3A). Examining the phospho-peptides that were quantified in all samples (i.e., no data imputation), we found that phospho-peptide intensities were strongly correlated in formalin-fixed and flash-frozen conditions (Fig. 3B). This data demonstrates that formalin-fixation does not alter phospho-proteomic identification or quantitation by LC-MS.

**Figure 3.**
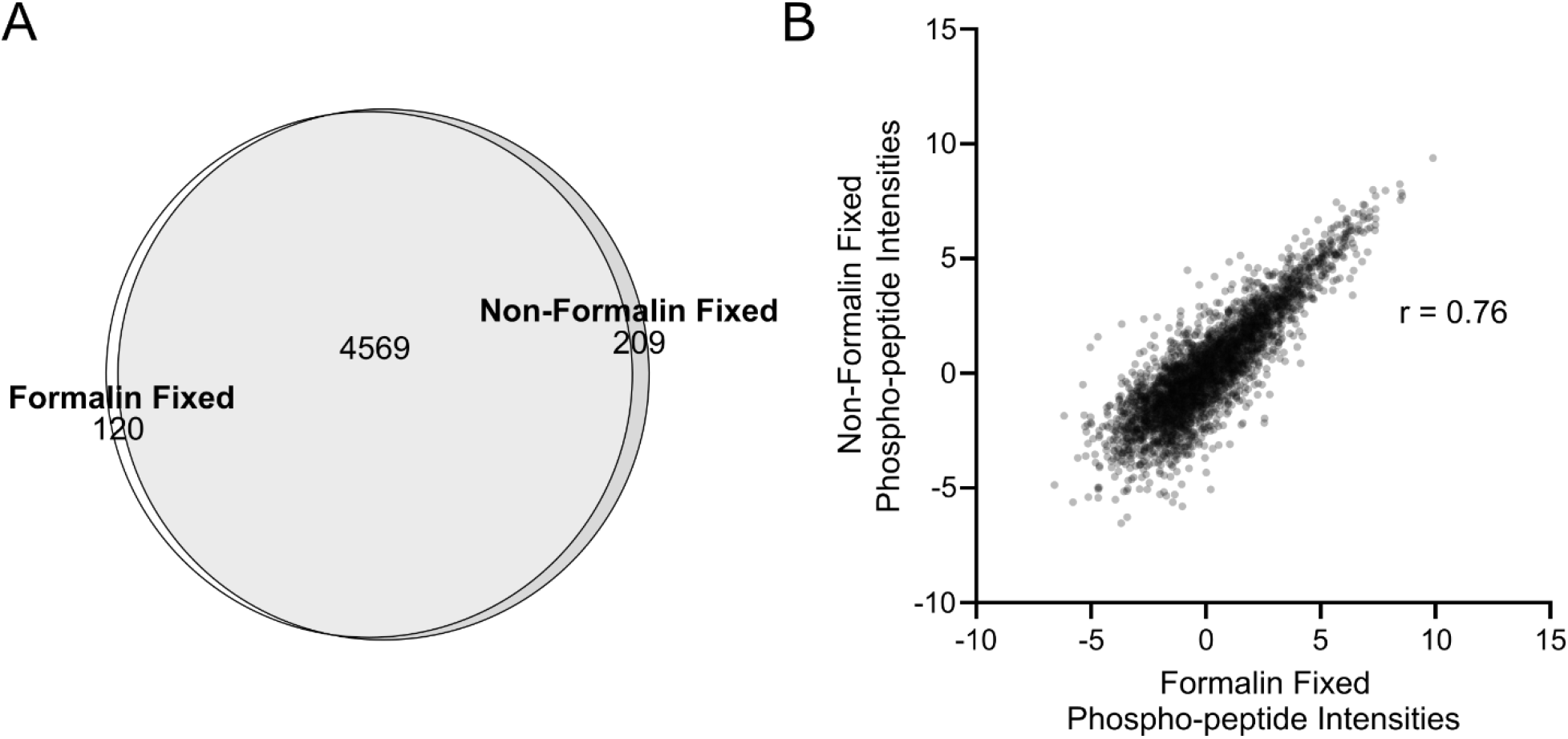
Formalin-fixation does not affect phospho-proteomics in T cells. A) The number of phospho-peptides identified by LC-MS in formalin fixed and non-formalin fixed samples is shown on a Venn diagram. Phospho-peptide recovery following fixation exceeded 93% in T cells. B) Phospho-peptide signal intensities were similar for formalin fixed and non-fixed samples (r = 0.76, p-value < 0.00001). Three independent biological replicates were used in each condition (Supp. Table 1).

### Magnet-activated cell sorting (MACS) enables purification of co-cultured CAR-T cells to >90%

Having demonstrated that formalin fixation does not affect phospho-proteomic data, we next tested whether we could isolate CAR-T cells from formalin-fixed co-cultures of CAR-T and SKOV3 cancer cells using magnet-activated cell sorting (MACS) (21). To assess the purity of CAR-T cell isolation, we first labeled SKOV3.CD19 and SKOV3.NT cells with the fluorescent marker CFSE. Then, we used flow cytometry to measure the percentage of CFSE^+^ SKOV3 cells during co-culture with anti-CD19 CAR-T cells and after MACS purification of CAR-T cells (Fig. 4A). After labeling, the percentage of CFSE^+^ SKOV3 cells reached 86.7-92.6% (Fig. 4B). Following mixing of SKOV3.CD19/NT cells at a 1:1 ratio with unlabeled anti-CD19 CAR-T cells, the percentage of CFSE^+^ cells dropped to 40.9-43.9%. Lastly, after formalin fixation and MACS purification, the percentage of CFSE^+^ cells was reduced to 5.8-7.6% of total live cells, supporting that the elution was >90% CAR-T cells. We also confirmed this MACS sorting efficiency using GFP-labeled SKOV3 cells (Supp. Fig. 1). Together, this data indicates that MACS can enrich CAR-T cells to high purity from formalin-fixed co-cultures of SKOV3 cancer cells and CAR-T cells.

**Figure 4.**
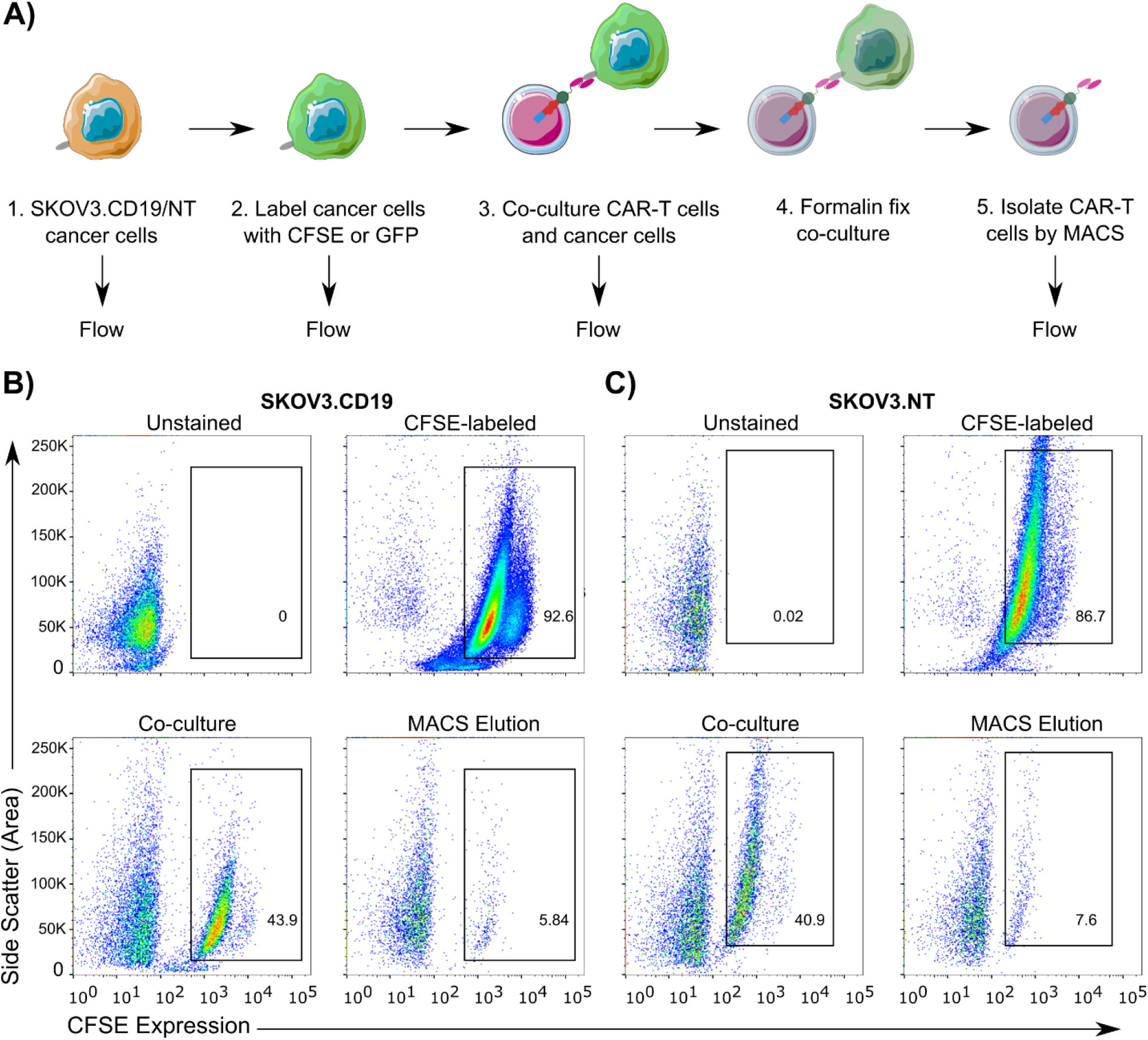
Assessing the purity of CAR-T cell isolation by MACS following co-culture with cancer cells. A) Workflow for assessing the purity of CAR-T isolation from formalin- fixed co-cultures of CAR-T cells with either SKOV3.CD19 or SKOV3.NT cells. B,C) Flow cytometry plots of SKOV3.CD19 (B) and SKOV3.NT (C) cells before labeling with CFSE (upper left), after labeling with CFSE (upper right), after 1:1 mixing with unlabeled CAR- T cells (lower left), and after MACS elution (lower right). Formalin fixation was performed after 45 min of co-culture. The purity of the isolated CAR-T cell eluates was >90%.

### Analysis of CAR-T cell signaling following stimulation by antigen-presenting cancer cells

Next, we applied our formalin fixation and MACS purification protocol to analyze CAR-T cell signaling. To measure activation of the CAR in response to its target antigen, we mixed anti-CD19 CAR-T cells with either SKOV3.CD19 or SKOV3.NT cells (i.e., negative control) at a 1:1 ratio. After 45 min, co-cultures were formalin fixed, and CAR-T cells were purified by MACS, followed by lysis, protein de-crosslinking, tryptic digestion, TiO2 phospho-peptide enrichment, and LC-MS proteomics. Across three biological replicates run in technical duplicate in each condition, we identified 4,141 phospho- peptides from 1,964 proteins, of which 2,327 phospho-peptides from 1,293 proteins were quantified in at least 10 of 12 samples and in at least one of technical replicate from each sample (Supp. Table 2). Comparing CAR-T cells stimulated with SKOV3.CD19 cells to CAR-T cells stimulated with SKOV3.NT cells, the log2 fold changes in phosphorylation across all phospho-peptides were normally distributed, with mean and median log2 fold change slightly greater than zero (Supp. Fig. 2). In addition, we found 171 phospho- peptides that were quantified in all six samples of SKOV3.CD19-stimulated cells but zero samples of SKOV3.NT-stimulated cells, suggesting that these phospho-peptides were strongly activated by CAR signaling. Conversely, we found only 2 phospho-peptides that were quantified in all six SKOV3.NT-stimulated samples but zero SKOV3.CD19- stimulated samples. Of the phospho-peptides found exclusively in SKOV3.CD19 stimulated CAR-T cells, three phospho-peptides were from MKI67 (pSer2105, pThr2692, and pSer538), a marker of cell proliferation (28), one was from EGFR (pSer991), and one was from MEK1/2 (pSer226/pSer222). Taken together, these data support that anti-CD19 CAR-T cells stimulated by CD19-expressing cancer cells exhibit upregulated signaling relative to cells stimulated by non CD19-expressing cancer cells.

We next compared our phospho-proteomic data to the known CAR signaling network (29). In our dataset, we quantified two phospho-sites on the CAR (CD3ζ pTyr142 and CD28 pTyr223) as well as 40 phospho-peptides from 33 proteins with known roles in CAR signaling including LAT, GRB2, SOS, RAF, MEK1/2, and ERK1/2 (Fig. 5). Notably, these phospho-peptides included signals thought to propagate through both the CD28 and CD3ζ activation domains (8, 11, 15, 30–32). Surprisingly, both CD3ζ pTyr142 and CD28 pTyr209 were slightly downregulated by CAR activation (log2 fold changes of -0.37 and -0.61, respectively). As expected, we observed activation of the MAPK pathway, including the activation sites of both ERK1/MAPK3 (pThr202/pTyr204, log2 fold change = 1.2) and ERK2/MAPK1 (pThr185/pTyr187, log2 fold change = 0.8). Interestingly, we observed >2-fold downregulation of pSer38 and pSer224 from LAT (linker for activation of T-cells family member 1), a protein which couples CAR activation to downstream kinases. Neither of these LAT phospho-sites has a known functional role, although Ser224 is close to Tyr220, a site whose phosphorylation regulates LAT activity (33). Lastly, we observed many upregulated phospho-sites on transcription factors including JUND pSer90 (log2 fold change = 2.6), JUN pSer63 (log2 fold change = 2.3), and ATF2 pSer112 (log2 fold change = 1.7). Together, this data show that our phospho-proteomic analysis quantified many signaling nodes in the known CAR signaling network, further validating the applicability of this approach.

**Figure 5.**
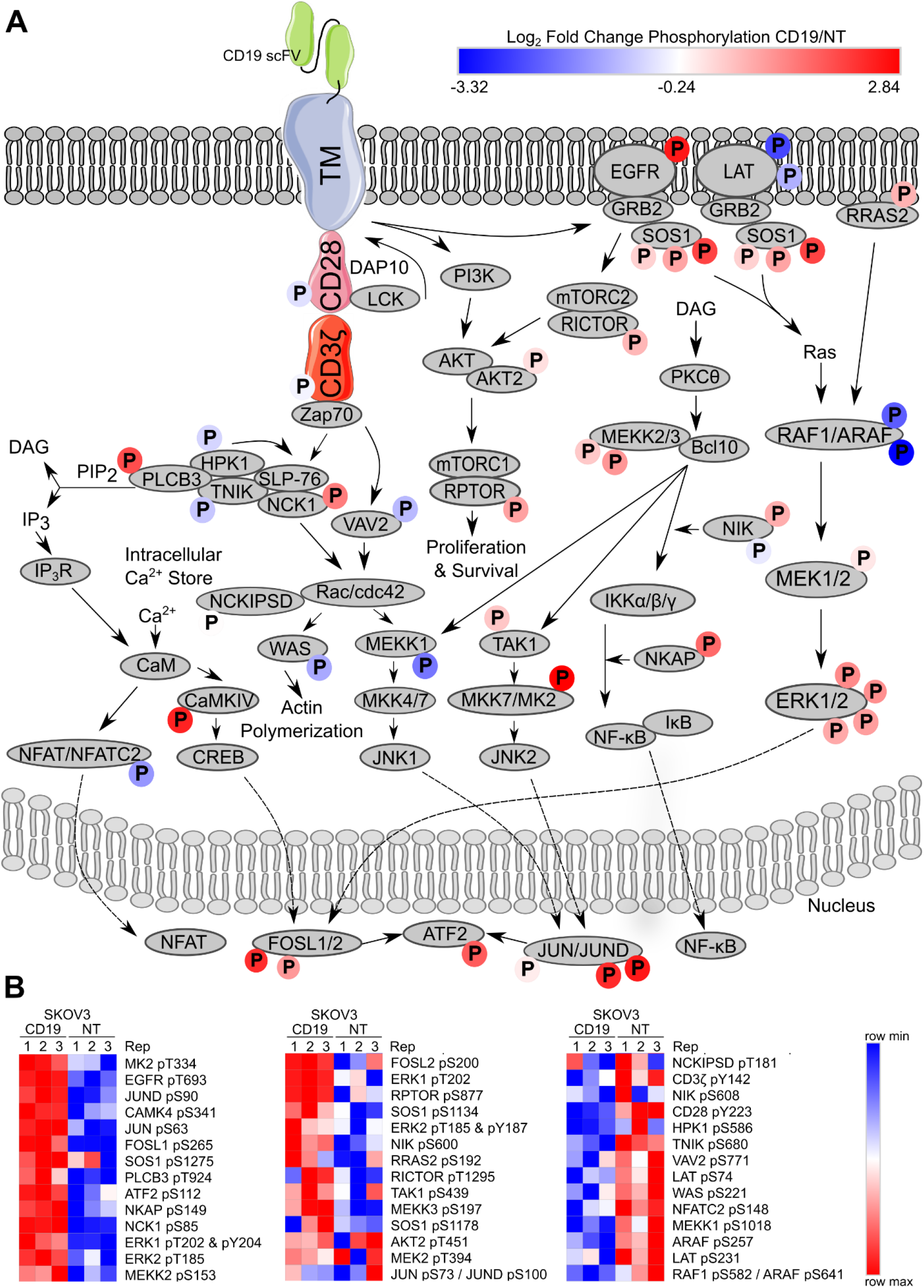
Phospho-proteomic analysis of the known CAR signaling network. CAR- T cells were stimulated for 45 min with cancer cells that either expressed CD19 (SKOV3.CD19) or not (SKOV3.NT). Co-cultures were formalin fixed, and CAR-T cells were isolated by MACS before phospho-proteomic analysis. Three biological replicates were analyzed in technical duplicate for each condition. A) Phospho-peptides identified in our dataset were matched to known CAR signaling pathways adapted from Cell Signaling Technology (29). Log2 fold change in phosphorylation levels of each phospho- peptide is indicated by the color of the circle enclosing a “P” according to the legend. B) Heatmap of individual replicate values of phospho-peptide levels for sites shown in A.

Next, to identify the most significant differences between CAR-T cells stimulated with SKOV3.CD19 cells and CAR-T cells stimulated with SKOV3.NT cells, we plotted our phospho-proteomic data on a volcano plot (Fig. 6A). The most significantly different phospho-peptides were also visualized on a heatmap to assess reproducibility (Fig. 6B). In total, 300 phospho-peptides were significantly upregulated and 67 were significantly downregulated in CAR-T cells stimulated with SKOV3.CD19 cells (FDR-adjusted p-value < 0.05). Notably, the significantly changing phosphorylation sites in our data set were significantly correlated with changes induced by antibody-coated bead stimulation of a similar CAR (Supp. Fig. 3) (11). We next annotated the significantly changing phospho- peptides with known regulatory roles from the PhosphoSitePlus database (34) This analysis identified phosphorylation sites including pSer251 of the ERBB receptor feedback inhibitor 1 (ERRFI1), a phosphorylation site that promotes EGF signaling by inhibition of the negative regulatory function of ERRFI1 (35), Thr693 of EGFR, whose phosphorylation can affect EGFR activity and trafficking (36, 37), and Ser17 of SRC, which induces enzymatic activity (38). Among the significantly downregulated phospho- peptides were Src-like-adapter (SLA) pSer190 and FYN-binding protein 1 (FYB) pSer56. Although neither of these phospho-sites has a known function, both SLA and FYB have known roles in T-cell receptor signaling (39–41). Taken together, these results highlighted both novel and known phosphorylation changes in anti-CD19 CAR-T cells stimulated by CD19-expressing SKOV3 cells.

**Figure 6:**
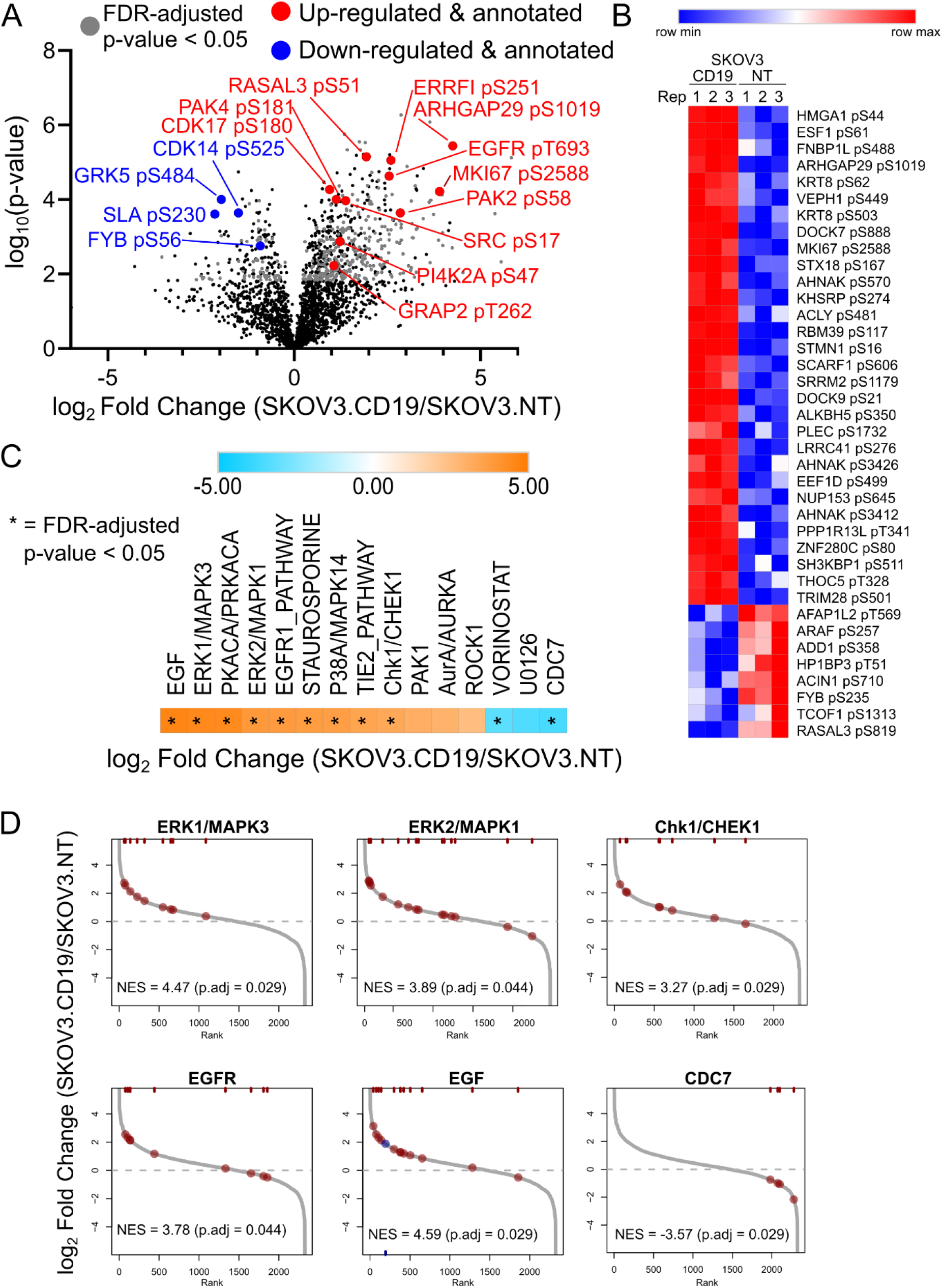
Analysis of CAR signaling following stimulation with antigen-expressing cancer cells. CAR-T cells were stimulated for 45 min with cancer cells that either expressed CD19 (SKOV3.CD19) or not (SKOV3.NT). Co-cultures were formalin fixed, and CAR-T cells were isolated by MACS before phospho-proteomic analysis. Three biological replicates were analyzed in technical duplicate for each condition. A) Phospho- proteomic data were plotted on a volcano plot to identify the most significant changes in phosphorylation levels. Phospho-peptides in grey represent FDR-adjusted p-value < 0.05 comparing SKOV3.CD19 stimulation with SKOV3.NT stimulation. Red (upregulated) and blue (downregulated) phospho-peptides have known regulatory roles according the PhosphoSite Plus database (34). B) The most significantly changing phospho-peptides (FDR adjusted p-value < 0.05 and log2 fold change >3 or <-2) were visualized on a heatmap to assess reproducibility. C) PTM-SEA identified significantly upregulated kinase, perturbation, and pathway signatures in CAR-T cells stimulated with CD19- expressing cancer cells. D) Rank-rank plots of significantly enriched kinase signatures identified by PTM-SEA.

Finally, to identify signaling pathways downstream of CAR activation, we analyzed our phospho-proteomic data using post-translational modification-signature enrichment analysis (PTM-SEA) (27). This analysis supported that CAR signaling activated the ERK1/MAPK3 and ERK2/MAPK1 signaling pathways (Fig. 5C,D), corroborating our observation of upregulated phosphorylation on the activation sites of these kinases. In addition, PTM-SEA identified upregulated phospho-signatures including EGFR, EGF, protein kinase A (PKACA), p38α/MAPK14, TIE2, and CHK1 as significantly upregulated in SKOV3.CD19-stimulated CAR-T cells. p38α signaling has also been observed in signaling downstream of a third-generation CAR, further corroborating a role of this kinase in CAR signaling (7). PTM-SEA also identified phospho-signatures of CDC7, a cyclin dependent kinase that regulates the cell cycle, and vorinostat, an inhibitor of histone deacetylases, as significantly downregulated in SKOV3.CD19-stimulated CAR-T cells. Taken together, these findings support that CARs activate both known and novel signaling pathways in CAR-T cells.

## Discussion

CAR immunotherapy has revolutionized cancer treatment. However, despite the rapid growth in clinical and preclinical research, the analysis of CAR signaling by phospho-proteomics (7, 9, 11, 15) and other methods (12, 13, 18, 42, 43) has been limited. In addition, current methods for phospho-proteomic analysis of CAR signaling require either bead-based antigen display to activate CAR signaling or stable isotope labeling (i.e., SILAC) to distinguish phospho-peptides from target and immune cells on the MS. Here, using formalin fixation and MACS, we have developed an efficient and cost-effective method for label-free phospho-proteomic analysis of CAR immune cells stimulated with antigen-presenting cancer cells. This method preserves transient phospho-signaling while recapitulating the physiological stimulus for CAR immune cells without the need for stable isotope labels. Using anti-CD19 CAR-T cells as an example, we have demonstrated the power of this method to reveal downstream CAR signaling that is consistent with findings from other phospho-proteomic approaches, and we expect that future applications will yield additional insight into how CARs encode the cytotoxic function of immune cells.

Because of the time requirement for MACS (∼2 h), formalin fixation is essential to ensure that CAR-mediated phosphorylation events, which occur on rapid time scales (44), are not lost during cell sorting. In proteomic analysis by phospho-flow, fixation with formalin or other agents is often used to preserve phosphorylation status (13, 45). However, for proteomic analysis by LC-MS, protein crosslinks must be reversed. The detergent SDS is ideal for reversing formalin crosslinks (and for solubilizing membrane proteins), but risks contaminating the LC-MS with a difficult to remove detergent. Here, we have taken advantage of the recent commercial development of suspension trapping (S-Trap) technology to remove SDS from cell lysates (25). Indeed, we found that formalin fixation and crosslink reversal resulted in no loss of phospho-proteomic information (Fig. 3), suggesting that this technique could be implemented more broadly for other rapidly changing post-translational modifications or protein-protein interactions.

Following formalin fixation, we used MACS to isolate CAR-T cells from SKOV3 cancer cells. Although fluorescence-activated cell sorting (FACS) can isolate immune cells for proteomic profiling (46), we found that FACS was unable to purify enough CAR-T cells for phospho-proteomics (e.g., 300 μg, not shown). MACS, in contrast, isolated CAR-T cells at >90% purity in an experimentally tractable amount of time (Fig. 4). Future work to improve our method will focus on increasing the efficiency of CAR-T isolation by MACS, potentially by using antibodies that more specifically recognize the CAR or by negative selection against cancer cell-specific markers. The ∼10% contamination of target cancer cells in our phospho-proteomic data represents a limitation compared to SILAC labeling of CAR-T and cancer cells (7, 15) or bead-based CAR activation (9, 11, 18) which analyze purer populations of CAR-T cells. However, the advantages of formalin fixation and MACS isolation in terms of cost, throughput, and realistic CAR activation by antigen-presenting cancer cells make this an attractive method for phospho-proteomic analysis of CAR signaling.

In our phospho-proteomic analysis, CAR stimulation with CD19-expressing SKOV3 cancer cells activated a broad range of signals (Figs. 5,6). Many of the observed signaling pathways are consistent with other analyses of CAR signaling, supporting the validity of our approach (7, 8, 11, 15). One of the strongest signals we observed was the activation of the ERK/MAPK signaling pathway, both at the level of individual ERK1/2 activation sites and PTM-SEA pathway analysis. Notably, The ERK/MAPK pathway plays a role in both CAR and T cell signaling, particularly in T cells that have not been primed with antigen (47–50). However, the functional implications of ERK signaling for downstream CAR-T function are currently unknown.

Beyond known CAR signaling pathways, our phospho-proteomic approach also identified novel signaling pathways downstream of the CAR. First, PTM-SEA identified CHK1, a serine/threonine protein kinase that regulates cell cycle processes (51), as significantly upregulated by CAR activation. CHK1 has primarily been investigated for its role in inhibiting replication stress in cancer cells (52), but it can also positively regulate EGF signaling in the absence of DNA damage by phosphorylating and inhibiting the negative regulator ERRFI1 on pSer251 (35). In our data, ERRF1 pSer251 was significantly upregulated by CAR activation (log2 fold change 2.60), supporting the possibility that CHK1 functions to modulate signaling in CAR-T cells. However, the role of CHK1 in CAR and TCR signaling has not yet been defined. In addition, our phospho- proteomic analysis identified that two EGFR phosphorylation sites with known regulatory roles (e.g., pThr693 and pSer991) (36, 37, 53) and an EGFR kinase signature were upregulated by CAR activation. Although a previous analysis of CAR-T cell signaling suggested that activated CAR-T cells could phosphorylate EGFR (43), it has traditionally been thought that hematopoietic cells do not express EGFR. However, sporadic reports have demonstrated that EGFR is indeed expressed by human Treg (54) and CD4^+^ T cells (55). Although we cannot exclude the possibility that these EGFR phospho-peptides come from SKOV3 and not CAR-T cells, proteomic profiling of pure CAR-T cells did support that CAR-T cells express EGFR (not shown). Thus, these data support the possibility that EGFR signaling may play an unappreciated role in CAR-T signaling. Our PTM-SEA also identified downregulation of phospho-signatures related to the kinase CDC7 and the histone deacetylase inhibitor vorinostat following CAR activation. Interestingly, in T cells, CDC7 activity may be required for T cell activation by affecting ERK and NFκB signaling but not proximal TCR signaling (56). Additionally, HDAC inhibitors have been shown to enhance the proliferation and survival of adoptively transferred T cells in cancer models (57, 58). As such, our results point to interesting CAR-T biology yet to be explored.

One limitation of our phospho-proteomic data is that we did not observe phosphorylation sites on several proteins known to be involved in CAR signaling including ZAP70, PLCγ1, and LCK. In addition, we did not capture some of the phosphorylation events known to occur on the co-stimulatory and activation domains of the CAR (i.e., CD28 and CD3ζ), including those present on the three immunoreceptor tyrosine-based activation motifs (ITAMs). Notably, many of these phosphorylation events occur on tyrosine residues (e.g., ITAM domains, ZAP70 pTyr492, LCK pTyr192, etc.), suggesting that inclusion of a parallel phospho-tyrosine enrichment (e.g., phospho-tyrosine antibody (15), Src SH2 superbinder (7)) would generate a more comprehensive view of the CAR phospho-proteome. Nevertheless, among the CAR phosphorylation sites that we did quantify, it is interesting that both were slightly downregulated (CD3ζ pTyr142 and CD28 pTyr209). This contrasts with previous phospho-proteomic studies where these phosphorylation sites were significantly upregulated by stimulation of second-generation CARs with antibody-coated beads (9, 11) or activation of a third-generation CAR with cell displayed-antigen (7). Potentially, this discrepancy could reflect differences in experimental design or that our CD19-expressing cancer cells have not maximally activated the CAR. Regardless, even in the absence of differential phosphorylation levels at these CAR sites, we observed robust activation of downstream signaling pathways (e.g., ERK) and target-cell killing. Further research will be necessary to clarify how differences in experimental design, CAR structure, and CAR activation methods regulate these important phosphorylation sties.

Taken together, our work represents the first label-free phospho-proteomic analysis of CAR signaling in CAR-T cells stimulated with antigen-presenting cancer cells. Our results provide insight into the signaling mechanisms downstream of CAR activation and suggest how CARs direct T cells to kill target cells. By coupling phospho-proteomic profiling with functional analysis of CAR-T cell cytotoxicity, we envision that an improved understanding of CAR signaling will enable engineering of next-generation CARs with increased potency, improved persistence, and reduced toxic side effects.

## Abbreviations

CAR: chimeric antigen receptor
CD: cluster of differentiation
CFSE: carboxyfluorescein succinimidyl ester
CHK1: checkpoint kinase 1
DTT: dithiothreitol
EGFR: epidermal growth factor receptor
EGF: epidermal growth factor
ERK: extracellular signal-regulated kinase
FA: formic acid
FACS: fluorescence-activated cell sorting
FF: formalin fixed
FYB: Fyn-binding protein
GFP: green fluorescent protein
IAA: iodacetamide
IFNγ: Interferon-gamma
ITAM: immunoreceptor tyrosine-based activation motifs
LC-MS: liquid chromatography-mass spectrometry
MACS: magnet-activated cell sorting
MAPK: mitogen-activated protein kinases
MEK: mitogen-activated protein kinase kinase
MEKK2: MEK/ERK kinase 2
MEKK3: MEK/ERK kinase 3
MK2: mitogen-activated protein kinase-activated protein kinase 2
NFF: non formalin fixed
PBMC: Peripheral blood mononuclear cell
PBS: phosphate buffered saline
PE: R-Phycoerythrin
PTM-SEA: post-translational modification-signature enrichment analysis
RT: room temperature
scFv: single-chain variable fragment
SKOV3.CD19: CD19-expressing SKOV3 cells
SKOV3.NT: non-CD19-expressing SKOV3 cells
SILAC: stable isotope labeling with amino acids in cell culture
SLA: Src-like adapter protein
SOS: Son of Sevenless
SOS1: SOS Ras/Rac Guanine Nucleotide Exchange Factor 1
TAK1: mitogen-activated protein kinase kinase kinase 7
TCR: T cell receptor
TCM: T cell media

## Acknowledgments

This work was supported by National Institute of Health grants (R01EB017206) and the USC Viterbi School of Engineering. Melanie A. MacMullan was partially supported by a scholarship from the ARCS Foundation. Some flow cytometric analysis was performed in the USC Flow Cytometry Facility that is supported in part by the National Cancer Institute Cancer Center Shared Grant award P30CA014089 and the USC Office of the Provost, Dean’s Development Funds, and the Keck School of Medicine of USC.

## Data availability

RAW LC-MS files and processed result files are available at MassIVE database (26) under identifier MSV000088632 (username: MSV000088632_reviewer, password: MM2021).

## Supplemental Figures

**Supplemental Figure 1.**
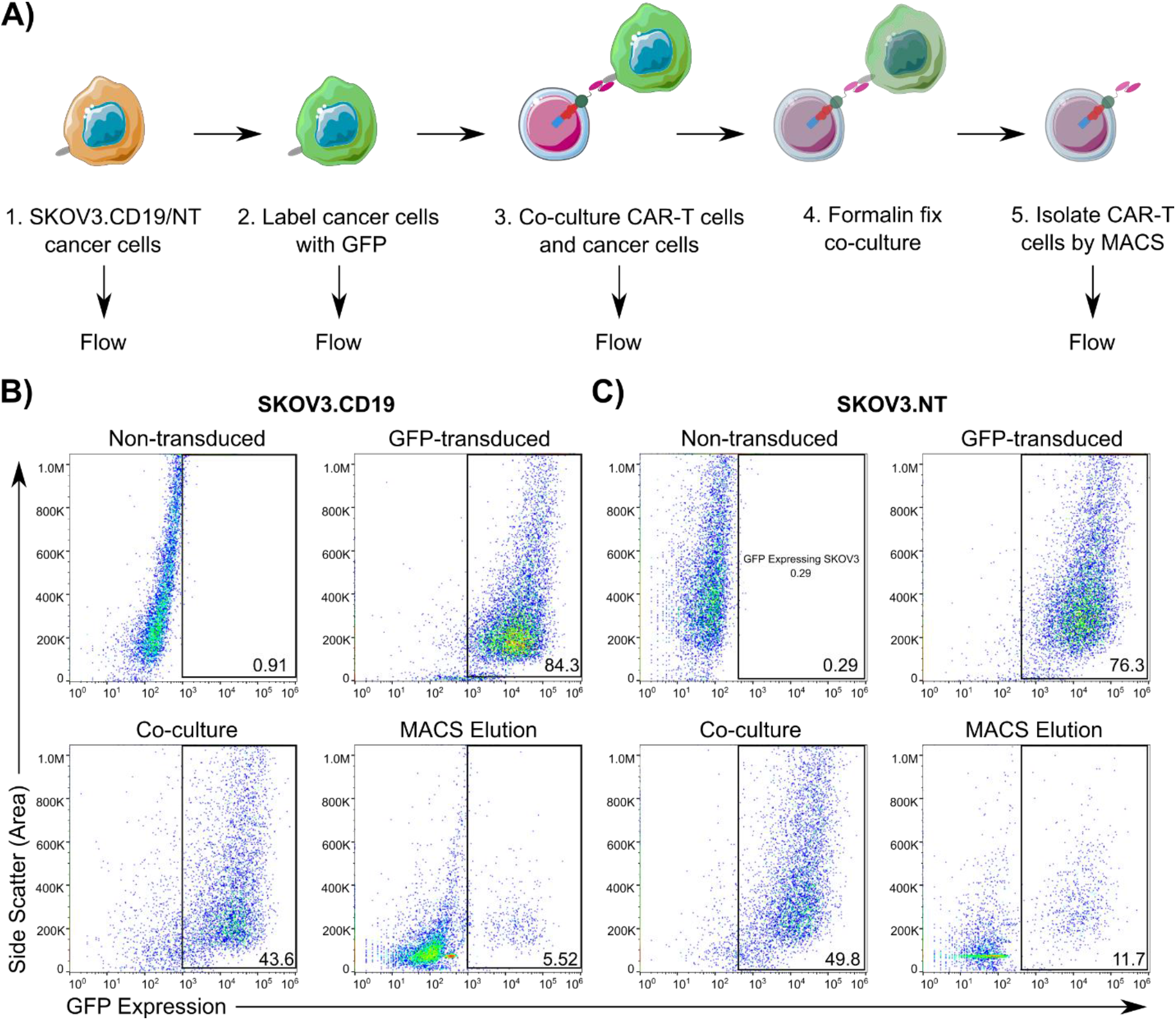
Assessing the purity of CAR-T cell isolation by MACS following co-culture with GFP-labeled cancer cells. A) Workflow for assessing the purity of CAR-T isolation from formalin-fixed co-cultures of CAR-T cells with either SKOV3.CD19 or SKOV3.NT cells. B,C) Flow cytometry plots of SKOV3.CD19 (B) and SKOV3.NT (C) cells before labeling with GFP (upper left), after labeling with GFP (upper right), after 1:1 mixing with unlabeled CAR-T cells (lower left), and after MACS elution (lower right). Formalin fixation was performed after 45 min of co-culture. The purity of the isolated CAR-T cell eluates was >90%.

**Supplemental Figure 2.**
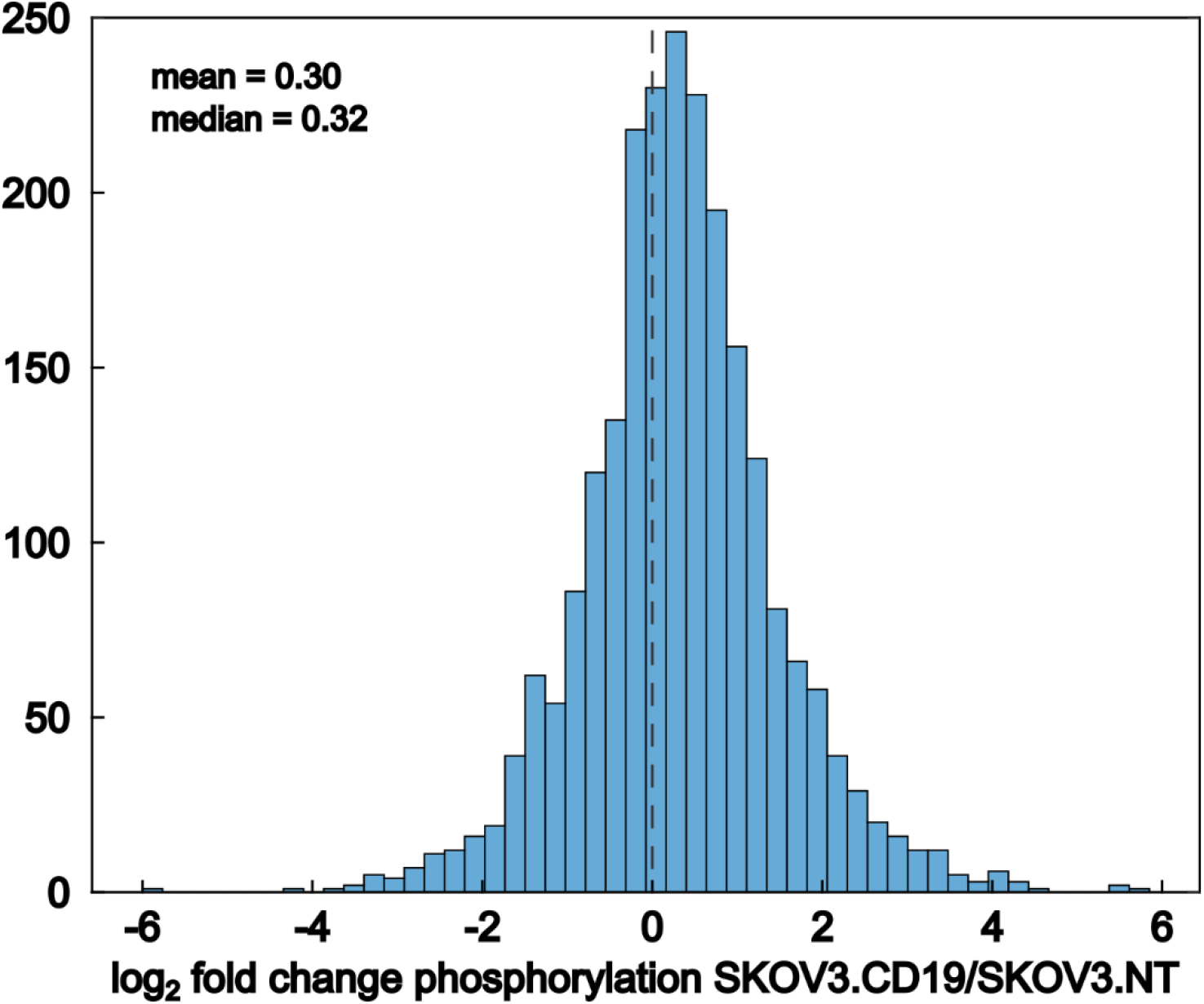
Histogram of log2 fold change in phosphorylation in CAR-T cells stimulated with SKOV3.CD19 or SKOV3.NT cells. Anti-CD19 CAR-T cells were mixed at a 1:1 ratio with either SKOV3.CD19 or SKOV3.NT cells. After 45 min, co-cultures were formalin fixed, and CAR-T cells were purified by MACS, followed by lysis and protein de-crosslinking, tryptic digestion, TiO2 phospho-peptide enrichment, and LC-MS proteomics (n = three biological replicates in each condition). The log2 fold change in phosphorylation comparing stimulation with SKOV3.CD19 to SKOV3.NT cells was calculated for 2,326 quantified phospho-peptides and plotted on a histogram.

**Supplemental Figure 3.**
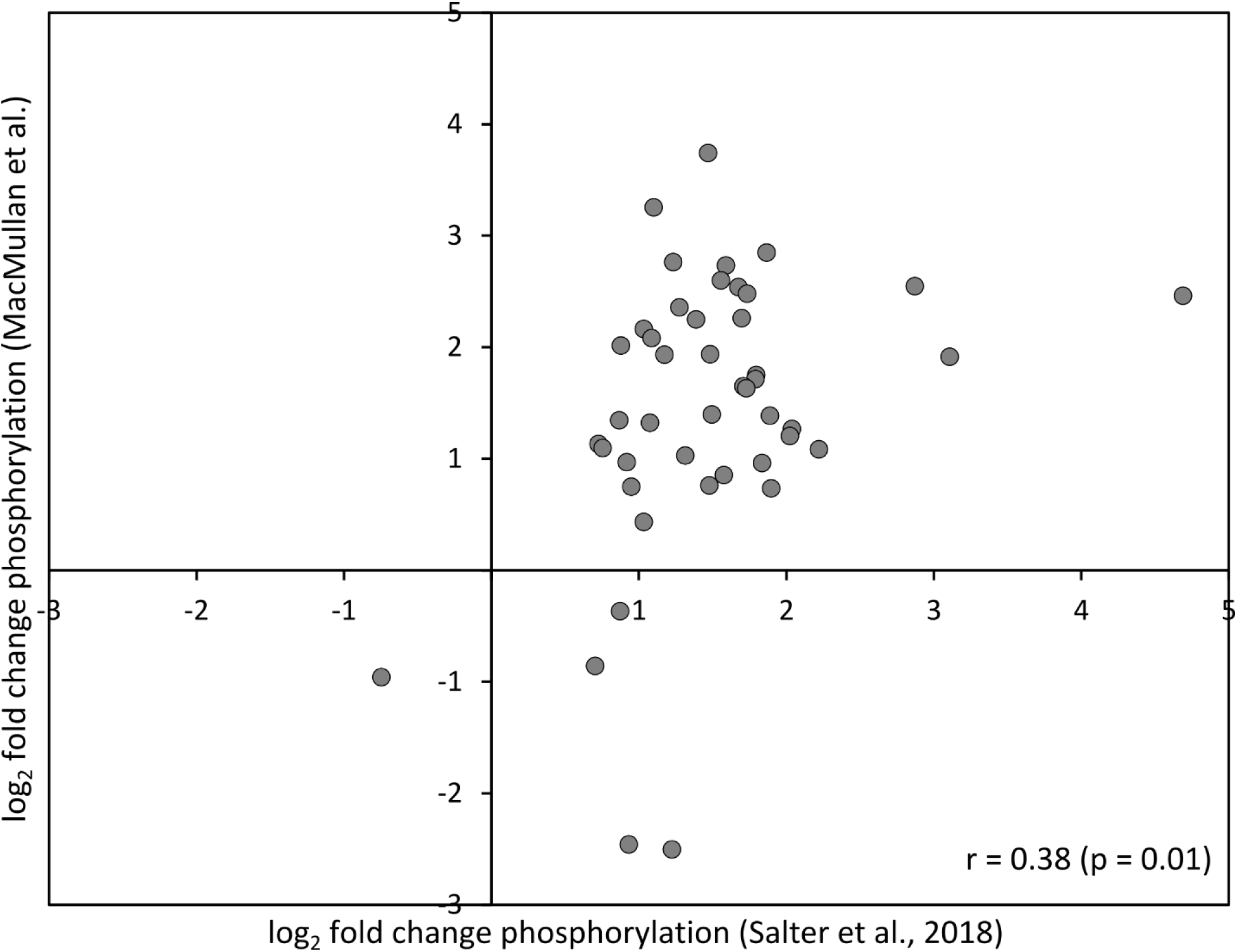
Comparison of significantly changing phosphorylation sites with Salter et al. Significantly changing phosphorylation sites from our data (Fig. 6, n = 367) were compared with significantly changing phosphorylation sites from Salter et al. (n = 1,189) (11). Both data sets were collected from T cells expressing a CD28/CD3ζ CAR after 45 min of stimulation with either CD19-expressing cancer cells (our data) or microbeads coated with an antibody that recognizes a Strep-tag II (STII) domain on the extracellular hinge of the CAR (Salter et al.). The overlap between the two data sets was significant (r = 0.38, n = 45, p = 0.01).

